# Tissue lamination state switches neuronal translocation from active to passive in the vertebrate retina

**DOI:** 10.64898/2026.07.02.736014

**Authors:** Margarida R. Cruz, Tiago Paixão, João Coelho, Caren Norden

## Abstract

Neuronal migration is essential for establishing functional tissue architecture in the developing central nervous system. While cell-intrinsic mechanisms driving neuronal movement have been identified, how migratory strategies adapt to dynamic developmental tissue changes remains less understood. Here, we address this question using retinal bipolar cells generated across unlaminated and laminated stages. This enables direct comparison of neuronal translocation across tissue states. Combining quantitative live imaging with targeted perturbations, we show that the migration mode of bipolar cells switches depending on tissue lamination state. Bipolar cells born before photoreceptor layer formation undergo directed, microtubule-dependent somal translocation. In contrast, later-born cells exhibit passive, non-directed displacement driven by collective tissue movements. Interference with tissue-wide movements impairs this displacement, while disrupting photoreceptor layer formation restores directed translocation. Independent of strategy, cells achieve accurate laminar positioning, indicating that tissue context determines neuronal migration mode, a principle likely relevant across the developing neural and other tissues.

## Introduction

In developing organisms, most cells are born at different locations than where they later function. Therefore, cell migration is a fundamental feature of embryogenesis and contributes to the formation, patterning, and organisation of most organs.

Correct cell migration is particularly important in the developing central nervous system (CNS) consisting of the brain, spinal cord and retina, where the precise positioning of neurons is a prerequisite for the formation of functional circuits.

Defects in neuronal migration can lead to diverse neurodevelopmental disorders, including retinal dysplasia, lissencephaly or microcephaly, and cortical dysplasia^1,2^. Many of the cellular and molecular mechanisms including guidance cues and signalling pathways, as well as the cytoskeletal components involved in cell movements^3–6^ have been explored, particularly in the developing neocortex. While these studies were insightful, the limited accessibility of cortical brain areas meant that cells were often examined outside of their native tissue environment. Furthermore, most investigations focused on the migrating cell itself^3,4,7^, meaning that the contribution of tissue context to neuronal migration is not as well explored. However, tissue context is highly relevant, as the developing brain is not a static scaffold but dynamically changes as development progresses. In early development, most brain areas consist of a pseudostratified neuroepithelium composed of one cell layer. During development, the neuroepithelium transforms into a highly laminated tissue hosting different types of neurons in distinct layers^8,9^. This means that neurons are exposed to changing cell composition and tissue architecture while development progresses. Additionally, it has been shown that neural tissues become more crowded as development proceeds^10^, which could also influence neuronal migration efficiency. Yet, whether and how neurons adapt their migratory behaviour to these progressive changes in tissue architecture remains poorly understood.

To address these questions, a system is needed that combines experimental accessibility and a well-characterised tissue architecture. Furthermore, it is important that neuronal movements can be studied in their native environment spanning the single cell as well as the tissue level. The vertebrate retina, particularly the zebrafish retina, fulfils these criteria. Its optical transparency, rapid external development, and its peripheral location make it ideally suited for live imaging.

The retina features a stereotypic arrangement of five neuronal cell types that is conserved across evolution: Retinal Ganglion Cells (RGCs) make up the basal GC layer, Photoreceptors (PRs) reside in the apical Outer Nuclear Layer (ONL), and interneurons, Amacrine Cells (ACs), Horizontal Cells (HCs) and Bipolar Cells (BCs), in the central Inner Nuclear Layer (INL) (Figure 1A)^8^.

**Figure 1.**
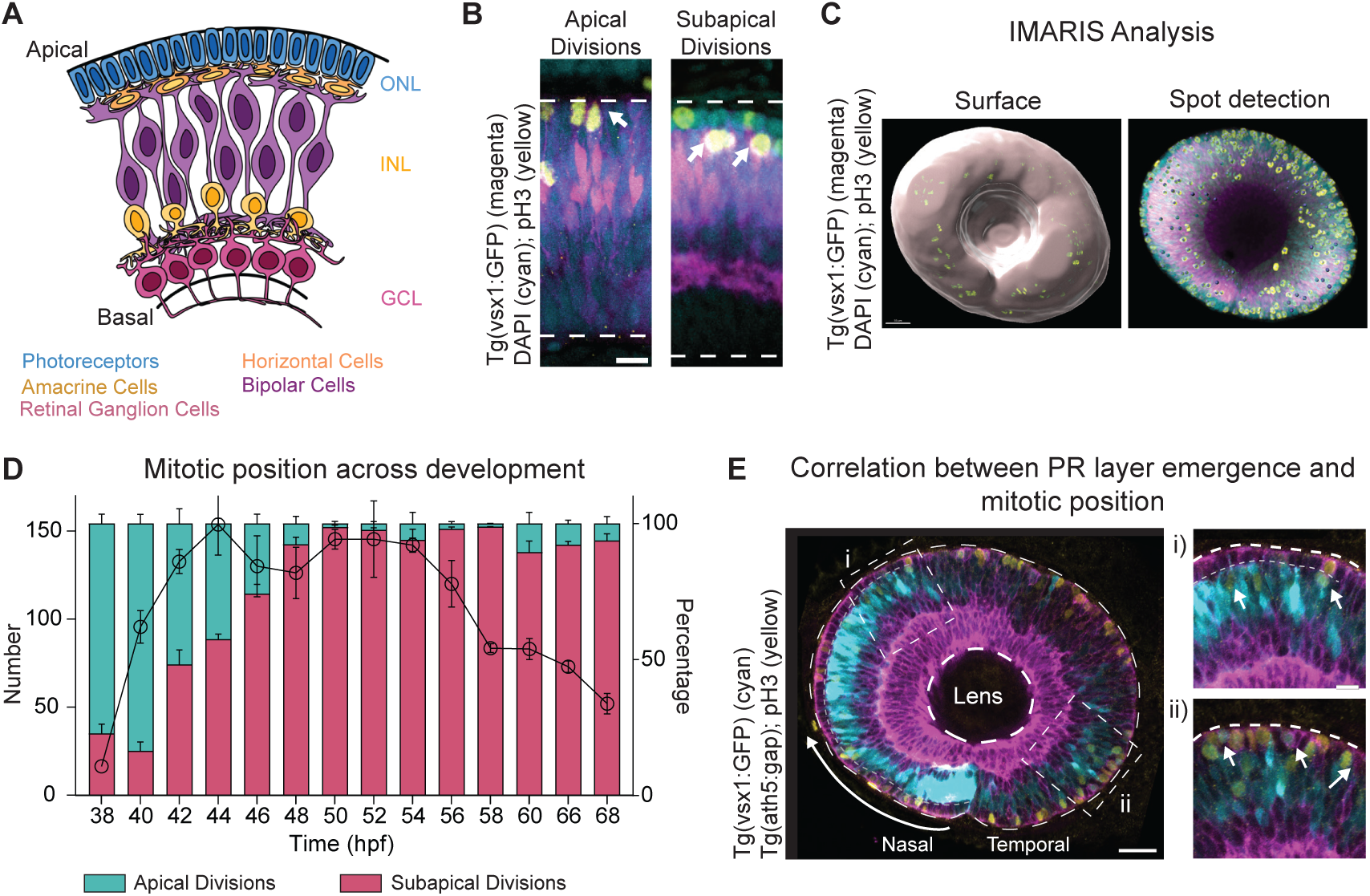
Mitotic position of Bipolar Cells depends on tissue lamination state. **(A)** Schematic representation of a cross-section of the zebrafish retina showing the final laminar position of the different retinal neurons. The cell bodies are organised into three layers from apical to basal: outer nuclear layer (ONL), inner nuclear layer (INL) and ganglion cell layer (GCL). Photoreceptors (PRs, blue) are located at the most apical layer (ONL). The interneurons, amacrine cells (light yellow), the horizontal cells (yellow) and the bipolar cells (BCs, purple) are distributed along the apico-basal axis of the INL. Retinal ganglion cells (pink) occupy the most basal layer (GCL). **(B)** Apical (left) and subapical divisions (right) at 40 hpf and 46 hpf, respectively. pH3 staining (yellow) labels dividing BC precursors (magenta). DAPI (cyan) staining shows tissue organisation and the presence of a PR layer. Arrows indicate divisions. Scale bar, 10 μm. **(C)** Three-dimensional surface reconstruction of a 48 hpf retina used for semi-automated quantification of mitotic BC position. Left: surface rendering used to define the apical surface boundary. Right: spot detection of pH3-positive (yellow) vsx1-positive (magenta) cells. Scale bar, 30 μm. **(D)** Quantification of total number of vsx1-positive cells (left axis) and percentage of apical (cyan) and subapical (magenta) vsx1-positive mitotic cells (right axis) across development (38 hpf – 68 hpf). Error bars represent mean ± SD. **(E)** Correlation between PR layer emergence and the shift in BC mitotic positions across development. pH3 staining (yellow) shows location of BC (cyan) mitosis in areas with PR (magenta) lamination and without PR lamination (i and ii respectively, magnified from white boxes, scale bar 10 μm). Arrows indicate divisions. Gray dotted line identifies PR layer. Differentiation wave progresses from nasal to temporal pole of the retina. Scale bar, 30 μm. *Multiply* plugin from Fiji was used to differentially decrease and increase Tg(*vsx1:GFP*) intensity in laminated and unlaminated areas, respectively.

All these retinal neurons are born at apical positions and must relocate to the sites at which they will later function. In the zebrafish retina, basal somal translocation is the predominant mode of neuronal migration at early developmental stages. In RGCs, PRs, and most likely also ACs and HCs, this movement depends on a stable, apically located, microtubule (MT) cytoskeleton^11,12^. Later in development, migration modes diversify. HCs and ACs switch to multipolar migration modes. This switch coincides with progressive changes in tissue architecture, suggesting an influence of tissue context on neuronal migration^8,13,14^. It was further shown that PRs undergo bidirectional movement that allows for the continued division of multipotent progenitors at the apical surface, thereby adapting their behaviour to the proliferative demands of the tissue^12^. Together, these examples show that tissue context is an important component of neuronal migration in the retina that deserves further attention and mechanistic dissection. Bipolar Cells (BCs) that are generated across an extended neurogenic window, spanning both unlaminated and laminated stages of retinal development, are an ideal model to ask how changing tissue environment influences neuronal movements as they can be followed along tissue lamination. Previous work^15,16^ revealed that BC precursors change their mitotic position dividing apically before PR layer formation and subapically upon PR layer emergence^12,16^. This indicates that progressive tissue layering influences their birth position. Whether and how changing birth position also influences translocation behaviour of BCs has, however, not yet been addressed.

To approach this question, we combined quantitative live imaging with cell and stage specific perturbations. We find that the apically born BCs undergo basal translocation with similar kinetics as RGCs and PRs and that, as in these cell types, this movement depends on a stabilised MT cytoskeleton.

This translocation mode was, however, not conserved in subapically born BCs, that displayed non-directed, MT-independent movements. This basal displacement was driven mainly by movements occurring in the surrounding tissue. Thus, the same neuron type can use different translocation strategies depending on place of birth and developmental stage of the tissue. Together, this data argues that tissue environment, alongside cell-intrinsic mechanisms, is a key determinant of neuronal migration strategies and needs to be considered to fully understand cell migration in the dynamically changing environment of the developing brain and beyond.

## Results

### Photoreceptor cell layer emergence determines Bipolar Cell Birth position

Bipolar Cells (BCs) can be born at apical and subapical positions and two studies suggested that mitotic position is linked to the developmental stage of the retina^15,16^. While both studies find that early born BCs are more likely to emerge apically than later born BCs, two not mutually exclusive hypotheses have been given to explain this observation: 1) the differentiation state of the surrounding post-mitotic BCs^15^; 2) physical hindrance by the emerging PR layer^16^. In the latter case, it was postulated that in the presence of a PR layer BCs can no longer reach the apical surface and therefore divide below PRs^16^. However, so far, the evidence was based on 2D optical sections which sampled only a small subset of mitotic events. To more quantitatively test for a link between PR layer emergence and BCs shifting to subapical birth positions, we characterised the spatio-temporal distribution of apically and subapically born BCs using the *Tg(vsx1:GFP)* transgenic line^17,18^. Vsx1 is a transcription factor that is initially expressed weakly in all progenitors and up-regulated in BC precursors and mature BC later in development^17,18^. We analysed 3-5 fixed embryos per stage at 2h intervals between 38 hours post fertilization (hpf), marking the onset of BC emergence, and 60 hpf, marking late stages of BC lamination. For later lamination stages, we analysed 3-5 embryos at 66 and 68 hpf. Mitotic cells were labelled by the mitotic marker phospho-histone H3 (pH3). pH3 Vsx1 double positive cells were analysed using IMARIS spot detection, defining apical divisions as divisions occurring within 10 µm from the apical surface, and subapical divisions as divisions occurring more than 10 µm away from apical surface, accounting for the thickness of the PR layer (Figure 1B-C).

Overall, the number of pH3 positive BCs increased from an average of 16.4 (± 6.4) at 38 hpf to 153.7 (± 30) at 44 hpf (Figure 1D). After 56 hpf, the number of pH3 positive BCs decreased to 120 (± 23) and 52 (±12.86) at 68 hpf.

When we analysed division positions, we noted that before 42 hpf, 85% (± 2%) of divisions occurred at the apical surface. From 42 hpf onwards, however, divisions occurred at both apical (52% ± 9%) and subapical positions (48% ± 9%). The proportion of subapical divisions peaked between 52 hpf and 54 hpf with 98% ± 0.5% of divisions occurring subapically (Figure 1D). Considering the whole developmental window analysed, 23% ± 8% occurred apically while 77% ± 8% of BC precursor divisions occurred subapically.

At 42 hpf, the timepoint at which BC divisions started to shift towards subapical positions has been shown to be the stage at which the PR layer starts to form^8^.

To more rigorously test this notion, we stained embryos from the *Tg(ath5:GAP-RFP;vsx1:GFP)* line against pH3 to follow BC division behaviour in the context of the forming PR layer marked by Ath5 expression (Figure 1E). In retinas at 42 hpf and 44 hpf, which featured both areas with an emerging apical PR layer and areas in which PRs did not yet occupy apical positions, apical BC divisions were seen only in PR-free regions while subapical divisions were seen upon PR layer formation (Figure 1E). This direct spatial correlation argued that indeed PR layer formation is the main driver for the shift in mitotic position of BC progenitors.

Together, our data shows that it is indeed the emergence of the apical PR layer that alters the division position of BCs.

### Apically born Bipolar Cells move basally with similar kinetics as seen for Retinal Ganglion Cells and Photoreceptors

Apically born BCs (aBCs) emerge mostly between 38 hpf and 42 hpf. This time window overlaps with the birth of neurons in the first neurogenic wave, RGCs, PRs, ACs, and HCs (Figure 2A)^8,11,12^. The translocation dynamics of these cell types have been studied previously and it was shown that, upon apical division, all these cells initially undergo directed basal somal translocation^11,12,14^. We thus asked whether these translocation dynamics are shared by basally moving aBCs. To investigate this, we mosaically labelled emerging BCs by either injecting a DNA construct containing a membrane marker, β-actin:mKate2-RAS, into the *Tg(vsx1:GFP)* line, or by transplanting cells from *Tg(vsx1:GFP)* embryos into WT embryos. BC behaviour after apical birth was followed between 38 hpf and 56 hpf at 5-minute intervals using light sheet microscopy due to its reduced phototoxicity^11,19^.

**Figure 2.**
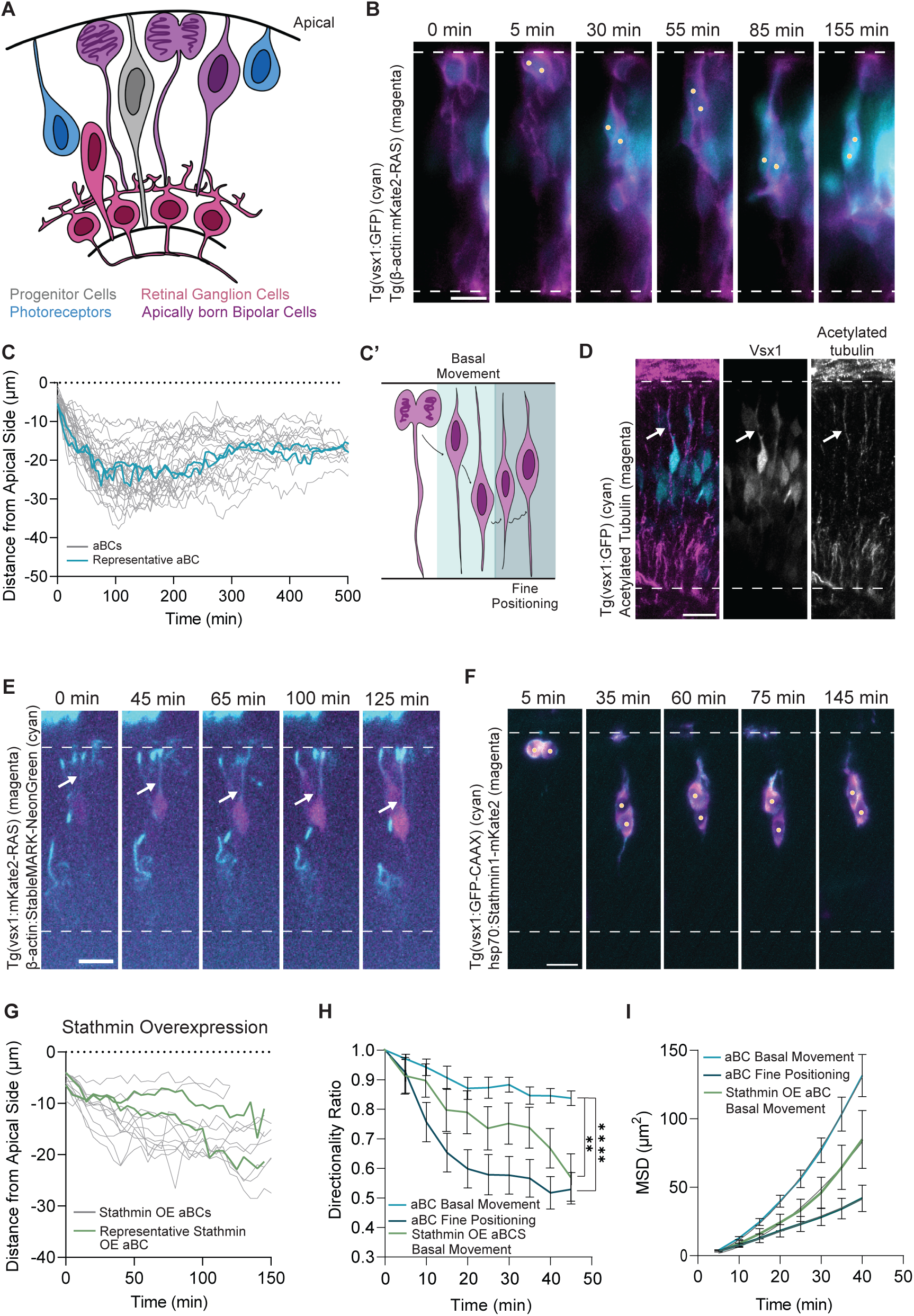
Apically born Bipolar Cells undergo MT-dependent somal translocation. **(A)** Schematic representation of the zebrafish retina illustrating the tissue context of apically born bipolar cells (aBCs, purple). Progenitor Cells (gray), PRs (blue), and RGCs (magenta) are translocating in earlier stages of BC neurogenesis. **(B)** Live imaging of aBCs translocation upon apical division. Time is shown in minutes. Yellow dots show BC sister cells followed. Dashed lines represent the apical surface and basal lamina. Scale bar, 10 μm. See Video S1 Part A. **(C)** Kinetics of BC translocation. Time 0 corresponds to the time of division. 48 single trajectories and a representative trajectory are shown. Dotted line marks the apical surface. (n=48 cells, N=17 embryos) **(C’)** Schematic illustrating the two phases of aBC translocation. Light blue phase corresponds to directionally persistent basal movement, and dark blue phase corresponds to fine positioning. **(D)** Distribution of acetylated tubulin (magenta) in early-born BCs (cyan) at 42 hpf. Arrows show enrichment of stable microtubules (MTs) along the apical process of BCs. Dashed lines represent the apical surface and the basal lamina. Scale bar, 10 μm. **(E)** Live imaging of stable MTs (cyan) in translocating aBCs (magenta). White arrows indicate the presence of stable MTs in the apical process of aBCs during basal translocation. Time is shown in minutes. Dashed lines represent the apical surface and the basal lamina. Scale bar, 10 μm. See Video S1 Part B. **(F)** Live imaging of translocating aBCs (cyan) following Stathmin overexpression (magenta). Dashed lines represent the apical surface and the basal lamina. Scale bar, 10 μm. See Video S1 Part C. **(G)** Kinetics of aBC translocation upon Stathmin OE. Time 0 corresponds to the time of division. 20 single trajectories and a representative trajectory are shown. Dotted line marks the apical surface. (n=20 cells, N=8 embryos) **(H)** Directionality Ratio of *Basal Movement* and *Fine Positioning* of unperturbed and Stathmin OE aBCs shown as the mean of all tracks ± s.e.m. (n=32 cells, N=16 embryos). P-values (Dunn’s Kruskal-Wallis Multiple Comparisons): 0.006 (aBC Basal Movement and Stathmin OE aBC Basal Movement); <0.0001 (aBC Basal Movement and aBC Fine Positioning). **(I)** Mean squared displacement (MSD) of Basal Movement and Fine Positioning of unperturbed and Stathmin OE aBCs shown as the mean of all tracks ± s.e.m. (n=32 cells, N=16 embryos).

Tracking 48 cells from 17 embryos revealed that after apical birth (Figure 2B and Video S1), aBCs translocated basally for an average of 126 minutes (± 27 minutes) moving on average 19 μm (± 6.3 μm) in the first 100 minutes (Figure 2C).

Following this initial basal movement, cells exhibited fine positioning within the INL (Figure 2C-C’ and Video S1). By modelling the translocation of each aBC through a Stochastic Differential Equation (see Methods), we were able to identify the time point that corresponded to the transition between *Basal Movement* and *Fine Positioning* for each cell. This allowed us to analyse and compare the kinetics of both phases of aBC movement (Figure 2H-I). Based on this data, Directionality Ratios of *Basal Movement* (ratio of 0.84 ± 0.14) were higher than for *Fine Positioning* (ratio of 0.53 ± 0.23) (Figure 2H). Analysis of Mean Squared Displacement (MSD) confirmed the notion that *Basal Movement* was more persistent and directional, as shown by the supralinear shape of the MSD curve with an α value of 1.66 ± 0.01, compared to an almost linear curve with an α value of 1.19 ± 0.02, indicating close to random movement (Figure 2I).

When comparing aBCs data with previously published data of RGC and PRs (acquired in our lab with similar microscopes and at the same time resolution^11,12^) we found that Directionality Ratios, MSDs, and instantaneous velocity values (Figure S1A) were very similar for all cell types (Table 1). Thus, retinal neurons born before PR layer emergence share kinetics of basal translocation with other apically born cells, namely PRs and RGCs.

**Table 1.** Summary of RGC, PR, and aBC kinetics during Basal Movement.

|  | **Retinal**<br>**Ganglion Cells**<br>**<sub>11</sub>** | **Photoreceptors**<br>**<sub>12</sub>** | **Apically born**<br>**Bipolar Cells** |
| --- | --- | --- | --- |
| *Duration* | 115 min | 95.5 min | 126 min |
| *Average displacement* | 28 $\mu\text{m}$ | NA | 19 $\mu\text{m}$ |
| *Average speed* | 14.9 $\mu\text{m}/\text{h}$ | 13.8 $\mu\text{m}/\text{h}$ | 17.4 $\mu\text{m}/\text{h}$ |
| *Median instantaneous velocity* | 0.26 $\mu\text{m}/\text{min}$ | NA | 0.18 $\mu\text{m}/\text{min}$ |
| *Final Directionality Ratio* | 0.88 | 0.73 | 0.84 |
| *MSD α-value* | 1.60 | 1.59 | 1.66 |

### Apically stabilised microtubules drive basal translocation of apically born Bipolar Cells

We next asked whether the similar kinetics seen for aBC translocation meant that also these cells depend on a stable apical MT cage as seen in RGCs and PRs^11,12^. We find that acetylated tubulin, a marker for stable MTs, was enriched in the apical process of aBCs in 40 hpf embryos (Figure 2D). To explore the dynamics of this stable apical MT cytoskeleton, we injected StableMARK^20^, a Kinesin-1 rigor construct that preferentially binds stable MTs, together with vsx1:GFP-CAAX to label BC membranes. Here, we used spinning disk microscopy for better subcellular resolution. Imaging sBCs between 40 hpf and 56 hpf at 5-minute intervals, showed that stable MTs became enriched in their apical process upon apical division. These MT bundles grew towards the basal side during basal aBC migration (Figure 2E). We never observed stable MTs in the basal process of BCs during translocation.

To understand whether these stable MTs were indeed driving basal aBC movement, we overexpressed the MT-destabilising protein Stathmin 1 under the control of the *hsp70* heat shock promoter, for temporal control^11,21^. Heat shock was applied at 36 hpf (39°C for 15 minutes) and light sheet imaging started 4 hours after heat shock (Figure 2F). 20 cells from 8 embryos showed that aBCs overexpressing Stathmin exhibited a shallower movement towards the basal side of the retina than what was seen for aBCs of controls (Figure 2F-G), moving on average 10.9 μm ± 6.6 μm, compared to 19 μm ± 6.3 μm (Figure S1C). This movement was significantly less directional than that of control aBCs (p-value=0.036; 0.56 ± 0.31 compared to 0.84 ± 0.14 in control embryos) (Figure 2H). Consequently, the MSD curve was flatter and showed a lower α value of 1.52 ± 0.04 compared to 1.66 ± 0.01 in control aBCs (Figure 2I). Our Stochastic Differentiation Model further validated the effect of Stathmin on the disruption of aBC basal translocation. When fitted with our Stathmin OE data, the model could not accurately predict the transition point (t_0_) from *Basal Movement* to *Fine Positioning* indicating that the force driving the movement towards the basal side was greatly reduced (Figure S1D).

These results show that, as in other apically born neuronal cell types, stable MTs are required for aBC basal movement. This suggests that all neurons born at early lamination stages show very similar translocation kinetics driven by the same mechanism, independently of cell identity and final position along the apicobasal axis in the mature retina.

### Subapically born Bipolar Cells move basally in a non-directed manner

While RGCs, PRs and aBCs maintain an apical process attachment during basal translocation, subapically born BCs (sBCs), retract their apical attachment prior to division and do not regrow it^16^ (Figure 3A, Figure 3C and Video S2).

**Figure 3.**
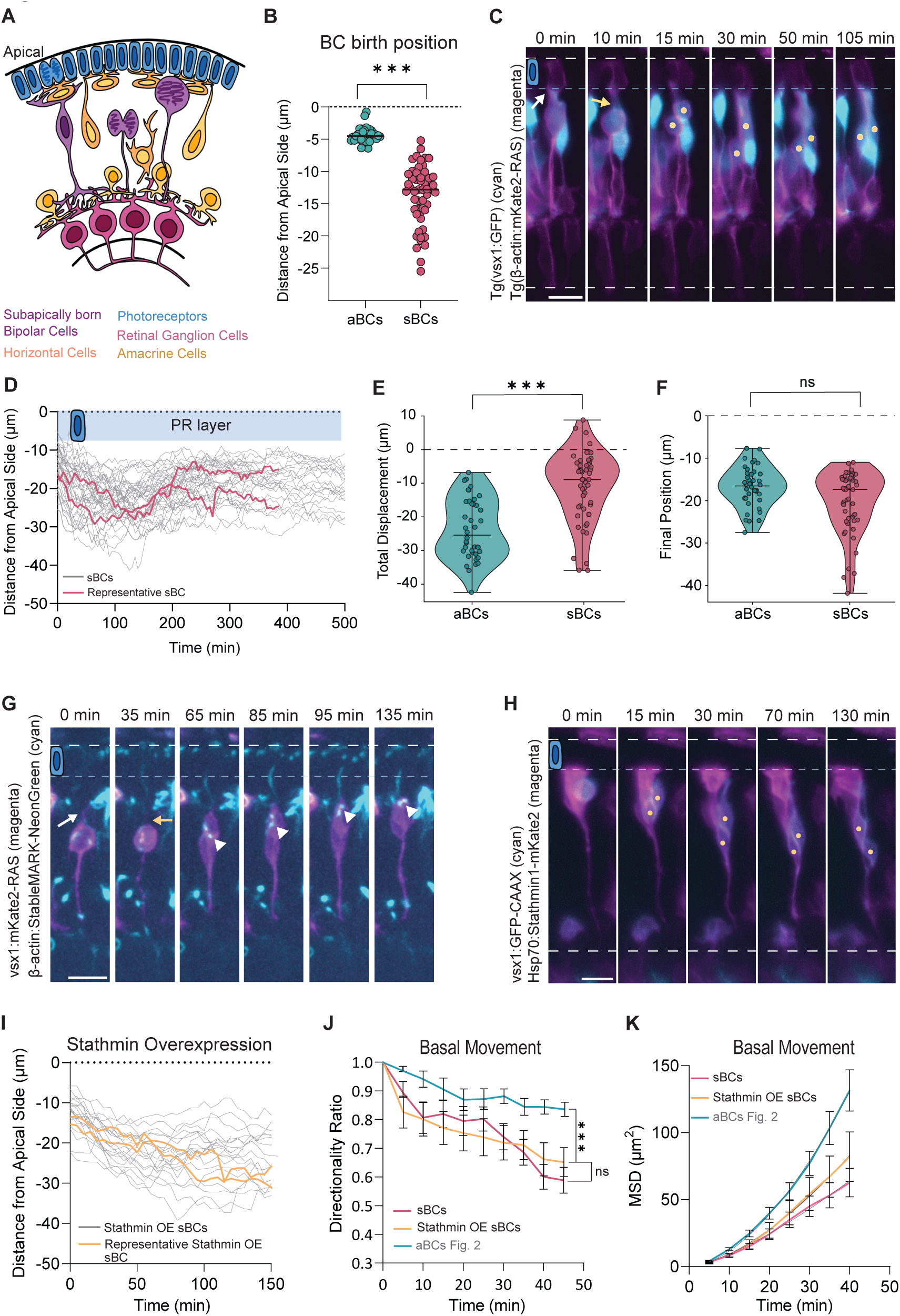
Subapically born Bipolar Cells undergo MT-independent translocation. **(A)** Schematic representation of a zebrafish retina illustrating the tissue context of subapically born bipolar cells (sBCs) (purple). PRs (blue) and RGCs (magenta) are already positioned in their respective layers. HCs and ACs are undergoing lamination, with some still migrating towards their final positions. **(B)** Distribution of BC birth position, measured as distance from the apical surface at division, for aBCs (cyan) and sBCs (magenta). Each dot represents one cell. Bar represents median value. P-value (Mann-Whitney Test) < 0.0001. **(C)** Live imaging of sBC translocation upon subapical division. Time is shown in minutes. Yellow dots show BC sister cells followed. White arrow points to the apical process. Yellow arrow points to retraction of the apical process. Dashed lines represent the apical surface and basal lamina. Scale bar, 10 μm. See Video S2 Part A. **(D)** Kinetics of sBC translocation. Time 0 corresponds to the time of division. 54 single trajectories and a representative trajectory are shown. Dotted line marks the apical surface. (n=54 cells, N=13 embryos). **(E)** Violin plot showing the distribution of the total displacement of aBCs and sBCs. Middle line is the median; whiskers are minimum and maximum; dots represent individual measurements. P-value (Mann-Whitney) < 0.001. **(F)** Violin plot showing the distribution of the final position of aBCs and sBCs. Middle line is the median; whiskers are minimum and maximum; dots represent individual measurements. P-value (Mann-Whitney)= 0.057. **(G)** Live imaging of stable MTs (cyan) in translocating sBCs (magenta). White arrowheads point to the centrosome in translocating sBCs. White arrow points to the apical process. Yellow arrow indicates the retraction of the apical process. Time is shown in minutes. Dashed lines represent the apical surface and the basal lamina. Scale bar, 10 μm. See Video S2 Part B. **(H)** Live imaging of translocating sBCs (cyan) following Stathmin overexpression (magenta). Dashed lines represent the apical surface and the basal lamina. Scale bar, 10 μm. See Video S2 Part C. **(I)** Kinetics of Stathmin OE in sBC translocation. Time 0 corresponds to the time of division. 22 single trajectories and a representative trajectory are shown. Dotted line marks the apical surface. (n=22 cells, N=9 embryos). (**J)** Directionality Ratio of *Basal Movement* of unperturbed and Stathmin OE sBCs shown as the mean of all tracks ± s.e.m. (n=38 cells, N=10 embryos). P-values (Dunn’s Kruskal-Wallis Multiple Comparisons): 0.0003 (sBC Basal Movement and aBC Basal Movement (see Figure 2H)); >0.9999 (sBC Basal Movement and Stathmin OE sBC Basal Movement). **(K)** Mean squared displacement (MSD) of *Basal Movement* and *Fine Positioning* of unperturbed and Stathmin OE sBCs shown as the mean of all tracks ± s.e.m. (n=38 cells, N=10 embryos).

We found that, as a result, sBCs show a much broader distribution of their birth positions along the apicobasal axis, ranging from 5.3 μm to 25.5 μm from the apical surface, compared to a maximum of 6.4 μm for aBCs (Figure 3B). Thus, given that some sBCs are born very basally, we asked whether active translocation is nevertheless needed in all these cells and whether the kinetics of this movement would be similar to aBCs.

To answer this question, we followed 54 sBCs from 13 embryos between 44 hpf and 68 hpf (Figure 3C and Video S2).

(Of note: When tracking and following fate decisions, we observed that a subset of these subapical divisions gave rise to asymmetric daughter cell pairs, producing one BC and one AC rather than two BCs (Figure S2A-B). This type of asymmetric division was not observed for apical BC divisions. Asymmetric BC divisions have been previously reported^22^, but at a higher frequency than observed here (41% as opposed to 17% observed in our dataset (Figure S2C)). This discrepancy could be due to the different labelling methods of BCs. Asymmetric divisions were excluded from our analysis.

Tracking symmetrically dividing sBCs, we found that these cells retract their apical attachment prior to division, as previously described (Figure 3C and Video S2)^16^. Upon division, sBCs move basally and afterwards exhibit a fine positioning phase, akin to what was seen for aBCs (Figure 3C-D). However, the basal movement was overall much shallower and only spanned 9 μm ± 7.1 μm from their respective birth position compared to 19 μm ± 6.3 μm for aBCs (Figure 3E). Instantaneous velocity analysis also showed a slower and more variable movement when compared to aBCs (Figure S1B). In line with this, Directionality Ratios were significantly lower at 0.59 ± 0.27 compared to 0.84 ± 0.14 for aBC (p-value= 0.0003; Figure 3J) and MSD analysis revealed an almost linear curve and an α value of 1.45 ± 0.02 (Figure 3K). Together, this argues that basal movements of sBCs are diffusive in nature differently to what was observed for aBCs.

To further confirm that aBCs and sBCs represent distinct translocation modes rather than cells that differ only on isolated parameters, we performed unsupervised PCA incorporating all kinetic parameters simultaneously (Figure S3A). The resulting clean separation of aBCs and sBCs in PCA space (p-value= 0.0001) shows that their kinetic differences are systematic and coordinated across all parameters, establishing two distinct translocation modes.

We conclude that BCs born after PR layer formation employ a different translocation mode with different, less efficient kinetics compared to earlier born aBCs.

### Subapically born Bipolar Cells do not rely on microtubules for basal translocation

While the diffusive kinetics of sBC argued against an MT driven mechanism, direct experimental confirmation was required. Acetylated tubulin staining showed no clear signal in sBC (data not shown). Further, live imaging of StableMARK labelled the centrosome but was not seen in other regions along the apicobasal axis (Figure 3G).

As done for aBCs, we overexpressed Stathmin at 42 hpf, the onset of sBC emergence (Figure 3H). No differences were observed in Directionality Ratio (0.65 ± 0.05) (Figure 3J) or MSD analysis (1.35 ± 0.52) upon Stathmin OE when compared to controls (Directionality ratio 0.59 ± 0.27, MSD 1.35 ± 0.36) (Figure 3K).

Thus, we concluded that sBCs translocation does not depend on stable apical MTs for basal translocation.

However, despite the different modes of translocation seen for aBCs and sBCs, both BC types reach similar laminar positions and the estimated final positions of the passively moving sBCs in our model were not significantly different from those of aBCs (Figure 3F). This suggests that active, MT-dependent movement is not a prerequisite for correct neuronal positioning.

### Subapically born BCs are passively displaced by collective tissue movements

Our results show that sBCs translocate in a non-directed manner, as shown by the MSD and Directionality Ratio measurements. Thus, we hypothesised that sBCs might be mainly displaced by other cell movements in the surrounding tissue, as previously seen for progenitor nuclei during IKNM which are passively moved basally by incoming nuclei after apical mitosis^23^. To test whether passive basal displacement could account for the translocating sBCs, we aimed to stall general tissue dynamics. To this end, we made use of the fact that stable MTs are responsible for efficient basal translocation of almost all retinal neurons, RGCs, PRs, and most likely also HCs and ACs^11,12^. We speculated that interference with basal movement in the majority of cells by Stathmin overexpression would reduce overall tissue dynamics at later lamination stages, when progenitor numbers are reduced and apical nuclear migration is negligible (Figure 4A).

**Figure 4.**
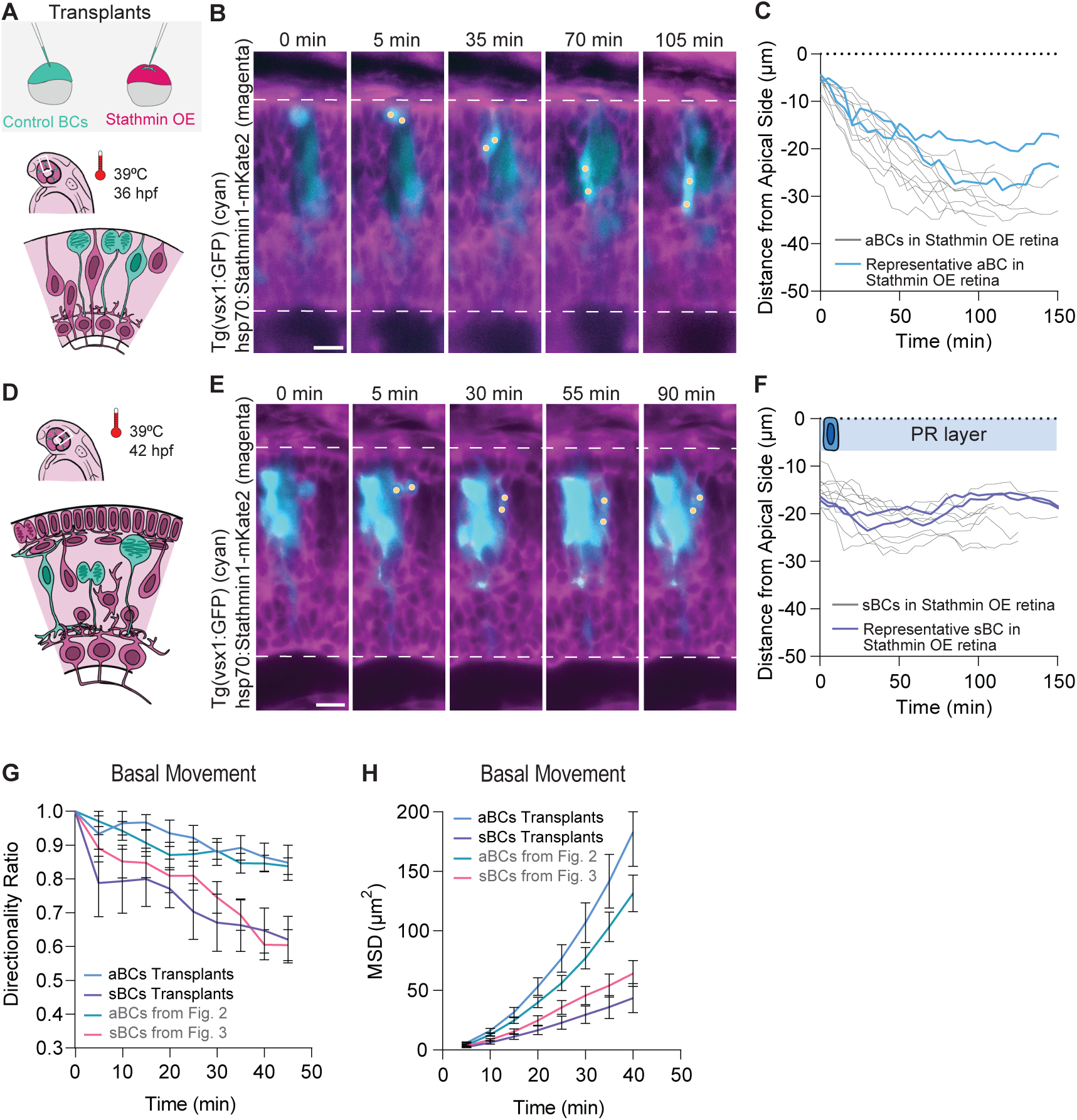
Subapically born Bipolar Cells translocation is dependent on collective tissue dynamics. **(A)** Schematic of the transplantation experimental design. Control cells Tg(*vsx1:GFP*) (cyan) from donor embryos are transplanted into host embryos Tg(*hsp70:Stathmin1:mkate2*) (magenta). To interfere with aBCs, heat-shock was performed for Stathmin overexpression at 36 hpf for 15 minutes at 39°C. **(B)** Live imaging of control aBC (cyan) translocation in a Stathmin OE host retina (magenta). Time is shown in minutes. Yellow dots show BC sister cells followed. Dashed lines represent the apical surface and basal lamina. Scale bar, 10 μm. See Video S3 Part A. **(C)** Kinetics of aBC translocation in Stathmin OE tissue. Time 0 corresponds to the time of division. 12 single trajectories and a representative trajectory are shown. Dotted line marks the apical surface. (n=12 cells, N=5 embryos). **(D)** Schematic of the experimental design. To interfere with sBCs, heat-shock was performed for Stathmin overexpression at 42 hpf, upon PR layer formation, for 15 minutes at 39°C. **(E)** Live imaging of control sBC (cyan) translocation in a Stathmin OE host retina (magenta). Time is shown in minutes. Yellow dots show BC sister cells followed. Dashed lines represent the apical surface and basal lamina. Scale bar, 10 μm. See Video S3 Part B. **(F)** Kinetics of sBC translocation in Stathmin OE tissue. Time 0 corresponds to the time of division. 14 single trajectories and a representative trajectory are shown. Dotted line marks the apical surface. (n=14 cells, N=4 embryos). **(G)** Directionality Ratio of *Basal Movement* of unperturbed aBCs (n=12 cells, N=5 embryos) and sBCs (n=14 cells, N=4 embryos) in Stathmin OE retinas shown as the mean of all tracks ± s.e.m.. **(H)** Mean squared displacement (MSD) of *Basal Movement* of unperturbed aBCs (n=12 cells, N=5 embryos) and sBCs (n=14 cells, N=4 embryos) in Stathmin OE retinas shown as the mean of all tracks ± s.e.m..

To study the movement of otherwise unperturbed cells, we transplanted cells from *Tg(vsx1:GFP)* control embryos into *Tg(Hsp70:Stathmin-mKate2)* embryos. Heat-shock was performed at 36 hpf to analyse aBC translocation (Figure 4A) and at 42 hpf to analyse sBC translocation (Figure 4D). In both cases, live imaging started 4 hours after heat shock.

Tracking of 12 transplanted aBCs from 5 Stathmin overexpressing embryos (Figure 4B-C and Video S3) showed that their basal movement was not affected by Stathmin overexpression in the surrounding tissue, as seen by very similar Directionality Ratios (Figure 4G) and MSD curves (Figure 4H) compared to unperturbed aBCs (final Directionality Ratio of 0.84 ± 0.14 and α-value of 1.66 ± 0.03). This was expected, as aBCs basal movement is active and cell-intrinsically driven by stable MTs.

In contrast, 14 transplanted sBCs in 4 Stathmin overexpressing embryos showed trajectories even flatter than in control environments (Figure 4E and Video S3), with cells moving on average 4 μm ± 3.8 μm compared to 9 μm ± 7.1 μm in control sBCs (Figure 4F). The Directionality Ratio showed even lower values (final Directionality Ratio of 0.59 ± 0.3) than those observed for sBCs in control environments (Figure 4G), and the MSD curve was even flatter (α-value of 1.40 ± 0.01) (Figure 4H).

This argues that sBCs are indeed passively displaced by the basal movement of the surrounding cells instead of actively translocating through the tissue as seen for aBCs. Thus, neuronal positioning can be achieved independently of active cell-intrinsic mechanisms.

### Disruption of PR layer integrity alters BC translocation behaviour at late developmental stages

When analysing sister cell behaviour upon division, we noted that aBC sister cells move basally in a highly correlated manner with almost overlapping trajectories (Figure 5A-B) and a Pearson correlation coefficient of 0.92 ± 0.10 (Figure 5C). In contrast, sBC sister cells movement is much less correlated. Most sBC sister cells instantly diverge from one another (Figure 5A-B) and show a correlation coefficient value of only 0.59 ± 0.31 (Figure 5C). This difference likely reflects the absence of an apical attachment after birth which most likely serves as the anchor for the MT-based force-generating mechanism that drives aBC basal displacement. If so, restoring apical attachment in late-born BCs after division should enable them to undergo directed translocation. To test this, we disrupted the PR layer after its formation by CRISPR-mediated depletion of the *prdm1b* transcription factor, which is involved in PR specification^24^. This treatment has been shown to induce ‘holes’ in the PR layer at stages when lamination is otherwise completed^25^. We confirmed the appearance of holes in the PR layer in Prdm1b-depleted embryos (Figure 5D) and used this condition to analyse late-born BC behaviour using the Tg(*ath5:GAP-RFP*) line combined with mosaically expressed vsx1:GFP-CAAX to label individual BCs (Figure 5C).

**Figure 5.**
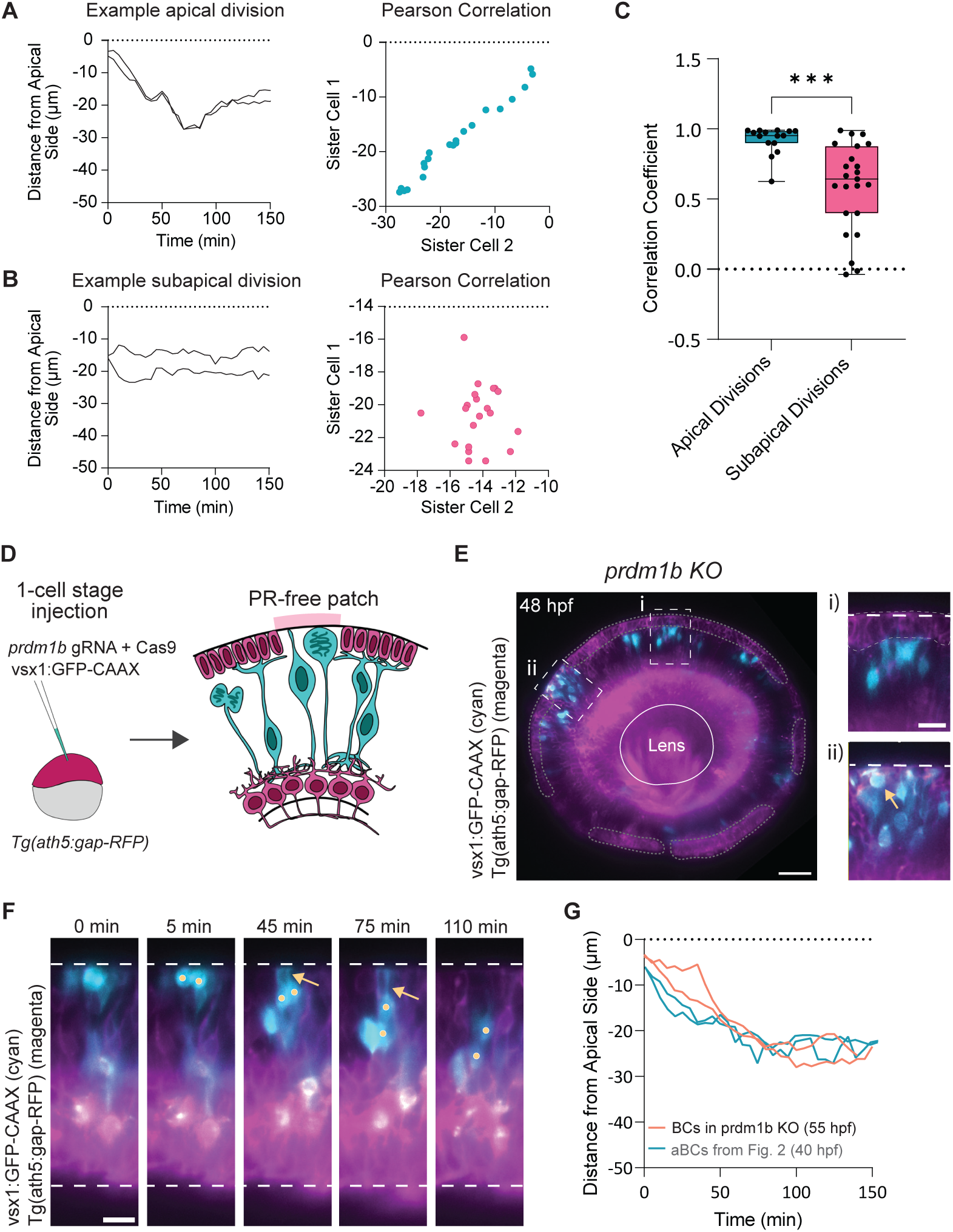
Disruption of PR layer integrity alters BC translocation behaviour at late developmental stages. **(A)** Kinetics of aBC sister cell translocation (left) and corresponding Pearson Correlation plot (right) to show the correlation between the movement of sister cells born at apical positions. **(B)** Kinetics of sBC sister cell translocation (left) and corresponding Pearson Correlation plot (right) to show the correlation between the movement of sister cells born at subapical positions. **(C)** Boxplot of the Pearson Correlation Coefficients for sister cell translocation from Apical Divisions and Subapical Divisions. Lower and upper box hinges are of 25th and 75^th^ percentiles, respectively; middle line represents the median; whiskers are minimum and maximum; dots represent individual measurements. Apical Divisions: n=16 sister cell pairs; Subapical Divisions: n=23 sister cell pairs. P-values (Kolmogorov-Smirnov Test) = 0.0003. **(D)** Schematic of the *prdm1b* KO experimental design. Embryos from the Tg(ath5:gap-RFP) line are co-injected with *prdm1b* gRNA + Cas9 complex and a vsx1:GFP-CAAX DNA construct at 1-cell stage generating embryos with interrupted PR layer (magenta) and mosaically labelled BCs (cyan). **(E)** Mosaically labelled BCs in a *prdm1b* KO; Tg(ath5:gap-RFP; magenta) embryo at 54 hpf. Areas with a more intact PR layer (dashed gray line; magnified area i) and areas with absent PR layer (magnified ii) are observed occupied by BC (cyan). Yellow arrow shows apical division of late-born BC in PR-depleted area (ii). Scale bar: 30 μm. **(F)** Live imaging of control late-born BC (cyan) translocation in PR-depleted area of a 56 hpf retina. Time is shown in minutes. Yellow dots show BC sister cells followed. Dashed lines represent the apical surface and basal lamina. Yellow arrow indicates the presence of an apical attachment. Scale bar, 10 μm. See Video S4. **(G)** Kinetics of late-born BC translocation in PR-depleted area upon apical division (orange line). Representative control aBC translocation (blue line). See Figure 2C. Time 0 corresponds to the time of division. Dotted line marks the apical surface.

Using light sheet live imaging starting at 48 hpf, when in controls the PR layer is complete and most BCs are born subapically, we were able to observe late-born BCs dividing within PR-depleted patches (Figure 5E and Video S4). While these events were rare, due to the stochastic nature of hole formation, cells dividing at the apical surface within these holes retained an apical attachment after mitosis and subsequently moved basally in a manner reminiscent of aBCs (Figure 5F-G and Video S4). Together, this argues that birth position, and the maintenance of the apical attachment, dictate the mode of translocation of BCs in the developing retina.

Together, this data demonstrates that apical attachment is both necessary and sufficient for directed BC translocation. Thus, it is the tissue lamination state, through its control of birth position and apical attachment, and not necessarily a cell-intrinsic program that determines translocation mode.

## Discussion

In this study we probed the influence of developmental stage and tissue environment on neuronal translocation using the retinal Bipolar Cells as a model. The extended neurogenic window of these cells, spanning unlaminated and laminated stages of retinogenesis, combined with the accessibility of the zebrafish retina, made them well suited to investigate their behaviour across lamination.

We revealed that depending on developmental stage and tissue lamination state, BCs implement different translocation strategies. While BCs born apically before PR layer formation use an MT-driven, cell-intrinsic mechanism, later born BCs dividing beyond the laminating PR layer undergo passive displacement driven by collective tissue movements. These findings show that, in addition to cell-intrinsic mechanisms, the tissue environment can be an important determinant of neuronal migration strategies.

A similar mechanism to the cell-intrinsic MT-driven migration mode of aBCs has already been observed for other retinal neurons, including RGCs and PRs. Taking previous data and data from this study into account, we show that all three cell types display similar kinetics and are dependent on stable apical MTs for translocation^11,12,14^. We speculate that a similar mechanism is also responsible for the initial basal translocation of ACs ^11^ and HCs. This would mean that MT-driven basal somal translocation is the default mechanism of early retinal neurons upon apical birth, before PR layer emergence. That this programme seems conserved between functionally and transcriptionally different cell types hints at a tissue-level rather than cell intrinsic origin of initial basal translocation.

It was previously shown that initial basal translocation of PRs, before they return to their final apical position, is crucial for the continued apical progenitor cell divisions, which is in turn important for the maintenance of tissue integrity^12,26^. This logic could extend to other neurons that are born during early retinogenesis, when proliferation and differentiation occur concurrently. Depending on cell type, this basal translocation can thus represent the primary driver of final positioning, serving mainly to clear the apical surface for ongoing proliferation. An important next step will be to investigate the molecular mechanism by which stable MTs drive basal somal translocation across neuronal cell types. As this movement mainly involves the basal translocation of the nucleus and cell soma, one possibility is that it is driven by physical coupling of the nucleus and the MT cytoskeleton via the LINC complex. Such a mechanism is seen in other contexts of nuclear translocation including the *Drosophila* eye disc, and the nuclear migration of *C. elegans* embryonic hypodermal cells^27–31^. Nuclear movements are directed towards the plus ends of MTs, so that this would most likely involve some type of kinesin motors. However, previous work from our group showed that perturbing the KASH proteins Syne1a and Syne2b or kinesins Kif1a and Kif5a did not affect basal translocation of RGCs^32^. Further, the fact that we do not observe MTs in the basal process of the cell argues against an efficient force generation by plus end directed transport. While together this makes a LINC-mediated process less likely, to formally exclude it will require systematic perturbation of the full spectrum of LINC complex components.

Another possibility for somal translocation across neuronal cell types is that the nucleus is actively pushed through MT plus end polymerisation, as seen for example for nuclear positioning in the Drosophila oocyte^33^. This would be consistent with a ratchet-like mechanism in which basally growing, and thereby pushing, MTs at the same time prevent the nucleus from returning apically which would in turn drive net basal movement. To test this idea, the StableMARK marker used here can be further exploited to visualise stable MT dynamics, in combination with nuclear envelope markers to detect possible MT-induced deformations. In addition, expansion microscopy could help to resolve MT-nuclear interactions.

In contrast to actively translocating aBCs, sBCs undergo passive, non-directed displacement that is driven by collective tissue movements. A key difference between the two BC cohorts is that aBCs inherit an apical attachment at birth while sBCs lose this attachment at division^16^. As the apical attachment hosts the centrosome that is anchored to the primary cilium, the organising centre for the stable MT bundles, it is most likely crucial for basal translocation^11,16,34^.

In sBCs, however, the centrosome becomes displaced once apical detachment is lost^16^. This most likely undermines a similar translocation mechanism as seen in aBCs. When later born BCs are allowed to divide apically in the *prdm1b* knock out condition, they retain their apical attachment and translocate basally with similar kinetics as aBCs. While we did not directly show that this reinstates MT-driven translocation, the data strongly argues that apical attachment is sufficient to initiate a cell intrinsic directed translocation mode regardless of developmental stage.

It is important to note that our results show that cell-intrinsic force generation, while prevalent across the developing CNS, is not an absolute prerequisite for correct neuronal positioning. At stages at which proliferation and lamination are largely complete, passive displacement by collective tissue movement can be sufficient to position newly born neurons correctly. It is possible that such passive displacement can only work over short distances as seen here for sBCs that only translocate tens of micrometres to reach their final position. Future studies will need to explore whether and under what circumstances passive displacement strategies of neuronal translocation extend beyond the retina in other brain areas, for example in the neocortex where late-born neurons must migrate past early born neurons to occupy more superficial layers^9,35^.

Our discovery of a passive translocation mode in the retina shows that the link between tissue architecture and neuronal translocation needs more exploration. One of the few examples that shows that the physical environment can influence neuronal migration modes was seen in a study on mouse cerebellar granule neurons. Combining *ex-vivo* and slice culture experiments, this study shows that actomyosin is enriched in the neurons’ leading process in 2D culture, generating a traction force that pulls the soma forward, but in 3D Matrigel or organotypic cerebellar slices actomyosin shifts to the posterior membrane, generating a pushing force, important for cell movements in the crowded tissues^36^.

Together with our study, this argues that the tissue environment could more broadly influence neuronal migration modes and their adaptation across the developing CNS and that more studies are needed that investigate this relationship.

Overall, extending studies of neuronal migration to the tissue perspective will help to explore the complexity of these phenomena. This in turn will have broad implications for the understanding of neurodevelopmental diseases. So far neuronal migration-associated disorders including focal cortical dysplasia or lissencephaly and microcephaly have predominantly been studied at the cellular level, with the main goal to identify cytoskeletal and other molecular players^37,38^. While this approach has been important in identifying causative genes of neurological conditions, it may not fully capture how disruptions to the broader tissue environment contribute to their emergence and severity. A better understanding of tissue-driven contributions to neuronal positioning could therefore reveal new pathogenic mechanisms and ultimately inform therapeutic strategies.

Current advances in live imaging, genetic tool development and in vivo reporters make it possible to address these questions in the native tissue context as seen in this study. This will be important to reveal how single cell dynamics and collective tissue behaviour interact during brain formation.

With the emergence of increasingly reproducible human retinal and other brain organoids, knowledge generated in tractable model systems like zebrafish, can be extrapolated to less accessible brain areas and human brain organoids. This will be crucial to more directly address the role of neuronal migration from cells to tissue during human brain development.

## Materials and Methods

### Zebrafish husbandry

Wild-type (*Danio rerio*; AB and Tupfel long-fin (TL) strains) and transgenic reporter lines were maintained and bred at 28°C (pH 7.0, conductivity 1000 μS/cm, 14:10 light:dark cycle; Techniplast). Embryos for experiments were raised in E3 medium at 28.5°C or 32°C. Staging of embryos was performed in hours post fertilization (hpf) according to Kimmel et al., 1995. Medium was replaced daily and supplemented with 0.2 mM 1-phenyl-2-thiourea (PTU, 10107703, Fisher Scientific) from 8 hpf onwards to prevent pigmentation. All zebrafish-related work was conducted in accordance with institutional standard operating procedures under the licencing of the DGAV (Direção Geral de Alimentação e Veterinária, Portugal) and in accordance with the European Union directive 2010/63/EU and with the Portuguese Decree Law n°113/2013.

### Zebrafish transgenic lines

To visualise emerging and mature Bipolar Cells and Photoreceptor Cells, *Tg(vsx1:GFP)*^18,40^, *Tg(ath5:gap-RFP*) ^41^, and double transgenic *Tg(ath5:gap-RFP, vsx1:GFP)* lines were used. To induce overexpression of Stathmin tissue-wide, the *Tg(Hsp70:Stathmin1-mKate2)*^12^ line was used. To visualise the membranes of cells, the *Tg(β-actin:mKate2-Ras)*^42^ line was used.

### DNA constructs and cloning

Gateway cloning (Thermo Fisher Scientific) based on the Tol2 Kit^43^ was used for all construct creation.

The plasmid with the vsx1 promoter sequence was a gift from Dr. Takeshi Yoshimatsu. To construct plasmids *vsx1:GFP-CAAX and vsx1:mKate2-Ras*, the vsx1 promoter sequence was PCR-amplified using Phusion Polymerase and primers forward (5’-GGGGGGGGACAACTTTGTATAGAAAAGTTGTATGGCAGTCAGTCAGCCCTTCTC-3’) and reverse (5’-GGGGGGGGACTGCTTTTTTGTACAAACTTGAGTCGATTCCGAACGAAGGGTAAAGTTTAAC-3’).

Gateway was used to recombine it with pME-EGFP-CAAX (Chien Lab, https://tol2kitkwan.genetics.utah.edu/index.php/PME-EGFPCAAX) or pME-mKate2-RAS ^11^ in the pTol2+pA R4-R2 backbone.

The sequence for StableMARK-mNeonGreen was obtained from plasmid pB80-hKif5b(1-560)rigor-2xmNeonGreen (a kind gift from Dr. Elias Barriga; plasmid 174649, Addgene;^20^). To construct plasmid *β-actin:StableMARK-mNeonGreen*, the StableMARK-mNeonGreen sequence was PCR-amplified using Phusion polymerase and primers forward (5’-GGGGACAAGTTTGTACAAAAAAGCAGGCTTAATGGCGGACCTGGCCGAG-3’) and reverse (5’-GGGGACCACTTTGTACAAGAAAGCTGGGTATTACTTGTACAGCTCGTCCATGCC-3’) and Gateway was used to recombine it with the *β-actin* promoter^43^ in the pTol2+pA R4-R2 backbone. To construct plasmid *β-actin:StableMARK-mKate2*, the StableMARK sequence was PCR-amplified using Phusion polymerase and primers forward (5’-GGGGACAAGTTTGTACAAAAAAGCAGGCTTAATGGCGGACCTGGCCGAG-3’) and reverse (5’- GGGGACCACTTTGTACAAGAAAGCTGGGTATTTTAGTAAAGATGCCATCATCTCAGCTGC-3’) and Gateway cloning was used to recombine it with the *β-actin* promoter and mKate2 (a gift from A. Oates Laboratory, École Polytechnique Fédérale de Lausanne (EPFL), Switzerland) in the pTol2+pA R4-R3 backbone.

To create plasmid *β-actin:DCX-mKate2*, the human Doublecortin (DCX) coding sequence was PCR-amplified from *β-actin:GFP-DCX*^11^ using Phusion polymerase and primers forward (5’-GGGGACAAGTTTGTACAAAAAAGCAGGCTTAATGGAACTTGATTTTGGACACTTTGACG-3’) and reverse (5’-GGGGACCACTTTGTACAAGAAAGCTGGGTACATGGAATCACCAAGCGAGTCC-3’) and Gateway was used to recombine it with mKate2 (a gift from A. Oates Laboratory, École Polytechnique Fédérale de Lausanne (EPFL), Switzerland) and the *β-actin* promoter in the pTol2+pA R4-R3 backbone.

Constructs *Hsp70:Stathmin1-mKate2* (plasmid 105969, Addgene; Taverna et al., 2016) and *Hsp70:mKate2-ras* (plasmid 105951, Addgene; ^34^) were published previously.

### DNA injections of zebrafish embryos

For mosaic labelling of cells in the zebrafish retina, purified plasmid DNA diluted in ddH_2_O supplemented with 0.05% of phenol red (P3532, Merck) was injected into the cytoplasm of one-cell stage embryos. Constructs were diluted from 10 to 30 ng/μL and did not exceed 40 ng/μL, even when multiple constructs were injected, and injection volumes ranged from 0.5-1 nL.

### Zebrafish F0 knockout of *prdm1b* by CRISPR-Cas9

To knockout *prdm1b*, embryos were injected with three crRNAs targeting asymmetrical exons 16, 22, and 51 of *prdm1b* as described in ^25^. The selected crRNAs were complexed with tracrRNA and Cas9, pooled and injected into the yolk of early 1-cell stage embryos as in ^45^ and ^25^. Exact injection protocol can be found in https://dx.doi.org/10.17504/protocols.io.5qpvo52wdl4o/v3.

### Blastomere Transplantations

To transplant blastomeres, donor and recipient embryos were placed in glass-bottom dishes and dechorionated at the 256-cell stage with 2 mg/mL Pronase (P8811, Sigma-Aldrich), followed by 4x rinse with Danieau’s Buffer (0.7 mM KCl, 58 mM NaCl, 0.6 mM Ca(NO_3_)_2_, 0.4 mM MgSO_4_, and 5 mM HEPES). Dechorionated embryos were transferred onto a 1 % agarose coated dish with 6×25 wells. At mid-blastula stage, 10-20 cells from donor embryos were transplanted into the recipient embryos using a micromanipulator equipped with a 0.1 mm diameter glass needle under an Olympus SZX10 stereo microscope. After transplantation, embryos recovered at 32 °C for 1-2 hours before being moved to 28.5 °C in agarose-coated dishes with fresh E3 medium supplemented with 0.2 mM PTU to prevent pigmentation.

### Heat shock

To induce the expression of the heat shock promoter (*hsp70*)-driven Stathmin1-mKate2, the petri dish containing the embryos was sealed and placed into a water bath set to 37°C or 39°C for 15-30 minutes. Screening and imaging of embryos started 2-3 hours after the heat shock.

### Whole-mount immunostaining

All immunostainings were performed on embryos fixed with 4% paraformaldehyde (PFA) (043368.9M, Thermo Fisher) in Phosphate-Buffered Saline (PBS), either 2-3 hours with agitation at room temperature (RT) or overnight at 4°C. After fixation, embryos were washed 4 x 15 minutes (min) with 0.5 % Triton-X in PBS (PBS-T 0.5%) followed by tissue permeabilization with 0.25% Trypsin-EDTA on ice for different periods of time depending on the developmental stage (10 min for 38 hpf, 15 min for 48 hpf - 58 hpf, and 18 min for 72 hpf - 120 hpf). Embryos were then rinsed 4x with PBS-T 0.8%. Blocking was performed in 10 % Normal Donkey Serum in PBS-T 0.8% for 2-3 hours at RT with agitation. Embryos were incubated with the following primary antibodies for 3 days at 4°C:

Anti-phospho-Histone H3 (Ser10) (06-570, Merck) 1:500;

Anti-acetylated α-tubulin (MABT868, Sigma-Aldrich) 1:200.

Embryos were washed five times for 30 min with PBS-T 0.8% and incubated for 2 days at 4°C with appropriate fluorescently labelled secondary antibody at 1:500.

The secondary antibodies used were the following:

Alexa Fluor 647 goat anti-rabbit (A-21245, Invitrogen)

Alexa Fluor 647 goat anti-mouse (A-21236, Invitrogen).

DAPI (MBD0015, Sigma-Aldrich) was added at 1:1000 to all experiments.

Embryos were washed four times for 15 min with PBS-T 0.8% and stored in PBS at 4°C until imaging.

### Image Acquisition

#### Laser scanning confocal microscopy

Fixed samples were mounted in glass-bottom 35mm dishes (P35G-1.5-14-C, MatTek) with 1% low-melting agarose (A9045, Sigma Aldrich). Samples were imaged using either a Leica Stellaris 5 upright confocal microscope or a Zeiss LSM 980 Airyscan2 inverted confocal microscope. The Leica Stellaris 5 upright confocal microscope was equipped with a HC PL APO 40x/1.1 Water Corr CS2 objective (#506425, Leica) and was operated by the LAS X (cv4.5.0.25531) software. The Zeiss LSM 980 Airyscan was equipped with a LD-C Apochromat 40x/1.1 Water Corr M27 water-immersion objective (Zeiss) and was operated using the Zen Blue v3.3 (Zeiss) software.

Acquired z-stacks spanned the entire thickness of the retina at 1μm step size.

#### Time-lapse imaging using spinning disk confocal microscopy

Live imaging of cytoskeleton dynamics in bipolar cells was performed using 5-10 embryos embedded into 0.6 % low-melting agarose (prepared in filtered E3 medium) in glass-bottom 35 mm dishes (P35G-1.5-14-C, MatTek). 15 minutes post embedding, when the agarose had solidified, the dish was filled with E3 medium containing 0.2 mM PTU and 0.04 % tricaine methanesulfonate (MS-222, 1004671, Pharmaq). Imaging was performed at 32°C using a Marianas SDC (Intelligent Imaging Innovations, 3i) spinning Disk confocal microscope, operated through Nikon Ti2 Eclipse and Slidebook 2024 (3i) software platform and equipped with Yokogawa CSU-W1 scanner plus the Uniformizer (Yokogawa), and Photometrics Prime 95B sCMOS (Teledyne) camera. Z-stacks of 80-100 μm at 1 μm step size, were acquired every 5 minutes using the 40x CFI Plan Apo Lambda S 40x Sil objective (Nikon). Embryos were imaged for a total of 10 to 16 hours.

#### Time-lapse imaging using LSFM

Live imaging of cell dynamics in zebrafish retinas was performed as previously described^11^ using the Zeiss Light Sheet Microscope operated with Zen 2014 SP1 software (v9.2.8.60) (black edition). In brief, the embryos were mounted in a 0.6% low-melting agarose (prepared in filtered E3) column. The sample chamber was filled with filtered E3 supplemented with 0.2 mM PTU and 0.04 % MS-222 maintained at 28.5°C. Entire retinas were imaged in a single-view at 5-minute time intervals for 10 to 24 hours. 120-160 μm z stacks were acquired with 1μm step size in a single view, dual sided illumination mode using the 10x/0.2 illumination objectives and a Plan-Apochromat 20x/1.0 W or 40x/1.0 W detection objective (all Carl Zeiss Microscopy) and the two PCO.Edge 4.2 sCMOS cameras.

### Image processing and analysis

All image data was processed in Fiji^46^. The raw LSFM data was first processed in Zen 2014 software (black edition, v9.2.8.54 and v9.3.1.393). Images were cropped, dual-side illuminators were fused or only one was selected using the Dual Side Fusion algorithm, and drift was corrected with a manual drift correction plugin (http://imagej.net/Manual_drift_correction_plugin, ImageJ). Image analysis of fixed samples was done using IMARIS (v10.0.0) and Fiji. Results were analysed and plotted using Microsoft Excel, GraphPad Prism version 9.4.0 for Windows (GraphPad Software), and Python.

The built-in *Multiply* plugin from Fiji was used to differentially decrease and increase Tg(vsx1:GFP) intensity in laminated and unlaminated areas, respectively in Figure 1E.

#### Cell tracking

Bipolar Cells were identified based on the expression of the vsx1 reporter and morphology. Cells were followed from birth in maximum projections of substacks on the time-lapse datasets using Fiji (https://imagej.net/plugins/mtrackj). Trajectories were obtained by tracing the centre of the cell body using the MtrackJ plugin in Fiji (https://imagej.net/plugins/mtrackj). Trajectories in y were offset to the apical surface at the time of division.

### Quantification and statistical analysis

#### Quantification of apical and subapical mitosis

DAPI staining of 38-72 hpf retinas was used to manually create a 3D retinal surface using the surface tool in IMARIS (v10.0.0). To quantify the number and percentage of pH3 vsx1 positive nuclei, the manually curated spots, obtained by the Spot Detection tool, were classified according to the *vsx1* channel intensity. Then spots were classified *apical* or *subapical* based on a 10 μm threshold from the retinal surface. Results were plotted in GraphPad Prism.

#### Analysis of migration kinetics of cellular movements

To calculate the instantaneous velocities, a custom-made Python 3 script was used that can be found here. MSDs and directionality ratios were calculated using the open-source computer program DiPer^47^, executed as a macro in Excel (Microsoft). To estimate directional persistence, the α-value was obtained by calculating the slope of the log-log plots of individual MSDs and time interval. To analyse the coordination between sister cells, a Pearson Correlation analysis was performed for the first 100 minutes of movement in Prism, and the Correlation Coefficients were extracted and compared between populations. For PCA analysis of the trajectories, a custom-made script was written using Python 3 which can be found here.

#### Statistics

Statistical tests were performed using GraphPad Prism v.9.4.0, Python and the DiPer program. Data was first tested for normality using D’agostino & Pearson and/or the Shapiro-Wilk test to decide between parametric and non-parametric tests. All statistical tests used are indicated in the figure legends. Likewise, exact *p*-values and sample sizes are provided in the respective figure legends.

### Statistical Modelling

We modelled cell position dynamics using a stochastic differential equation (SDE) framework that captures both deterministic movement trends and random fluctuations. The model assumes cells undergo a characteristic shift in their movement behaviour at a specific time point, which we refer to as the “switch time.” This is modelled using a sigmoidal transition function that smoothly interpolates between an initial movement regime (baseline drift) and a final regime (switched state).

The SDE includes: (1) a deterministic drift component describing the average movement tendency, (2) a damping term that brings the system toward an equilibrium position, and (3) a stochastic noise term accounting for biological variability. We used a hierarchical Bayesian approach where each cell trajectory has individual parameters, drawn from population-level distributions that are estimated from the data.

#### Inference and Model Fitting

Model parameters were estimated using Bayesian inference with the No-U-Turn Sampler (NUTS), a variant of Hamiltonian Monte Carlo, using Python package pymc v3.11. Priors were chosen to be weakly informative: exponential distributions for rate parameters and gamma distributions for positive-valued parameters. The model was fit separately to each of the experimental groups (WT-Apical, WT-Subapical, Stathmin-Apical, Stathmin-Subapical). Posterior distributions were summarized to obtain parameter estimates and credible intervals, and model fits were validated by simulating trajectories from the estimated parameters and comparing them to observed data.

### Declaration of generative AI and AI-assisted technologies in the writing process

During the preparation of this work, the authors used GPT-5.2 and Claude 4.6 to write and debug code and to edit the manuscript for spelling, grammar, and clarity. The authors reviewed and edited the output of these tools as needed and take full responsibility for the entire content of the publication.

## Acknowledgments

We thank all members of the whole Norden laboratory for lively and fruitful project discussion. Renata Cunha gave cloning and experimental support. We thank Diana Garcia-Morales, Ana Patricia Ramos, and Mariana Maia-Gil for valuable project discussion and Catarina Figueiredo for thoughtful feedback on the manuscript and experimental support. We are grateful to the Advanced Imaging and Aquatic facilities at the Gulbenkian Institute for Molecular Medicine (GIMM, formerly IGC). We are grateful to Dr. Elias Barriga for gifting us with the β-actin:StableMARK-2xNeonGreen plasmid. Funding: M.R.C. was a member of the integrative Biology and Biomedicine PhD program and was supported by the Fundação para a Ciência e a Tecnologia PhD studentship (UI/BD/152256/2021). T.P. was supported by the Fundação Calouste Gulbenkian-IGC, and by the Fundação para a Ciência e a Tecnologia CEEC (CEECINST/00085/2021/CP2984/CT0004). C.N. was supported by the Fundação Calouste Gulbenkian-IGC, by the European Research Council (ERC) under the European Union’s Horizon 2020 research and innovation program (grant agreement no. 819046), and by the Fundação para a Ciência e a Tecnologia CEEC (2023.07063.CEECIND/CP2854/CT0002).

## Author Contributions

M.R.C.: Conceptualisation, data acquisition and curation, investigation, methodology, validation, visualisation, formal analysis, writing (original draft). T.P.: methodology and data curation regarding model, writing (editing). J.C.: methodology concerning DNA constructs and line creation, writing (editing). C.N.: Conceptualisation, supervision, project administration, funding acquisition, writing (original draft).

## Supplemental Material

**Figure S1.**
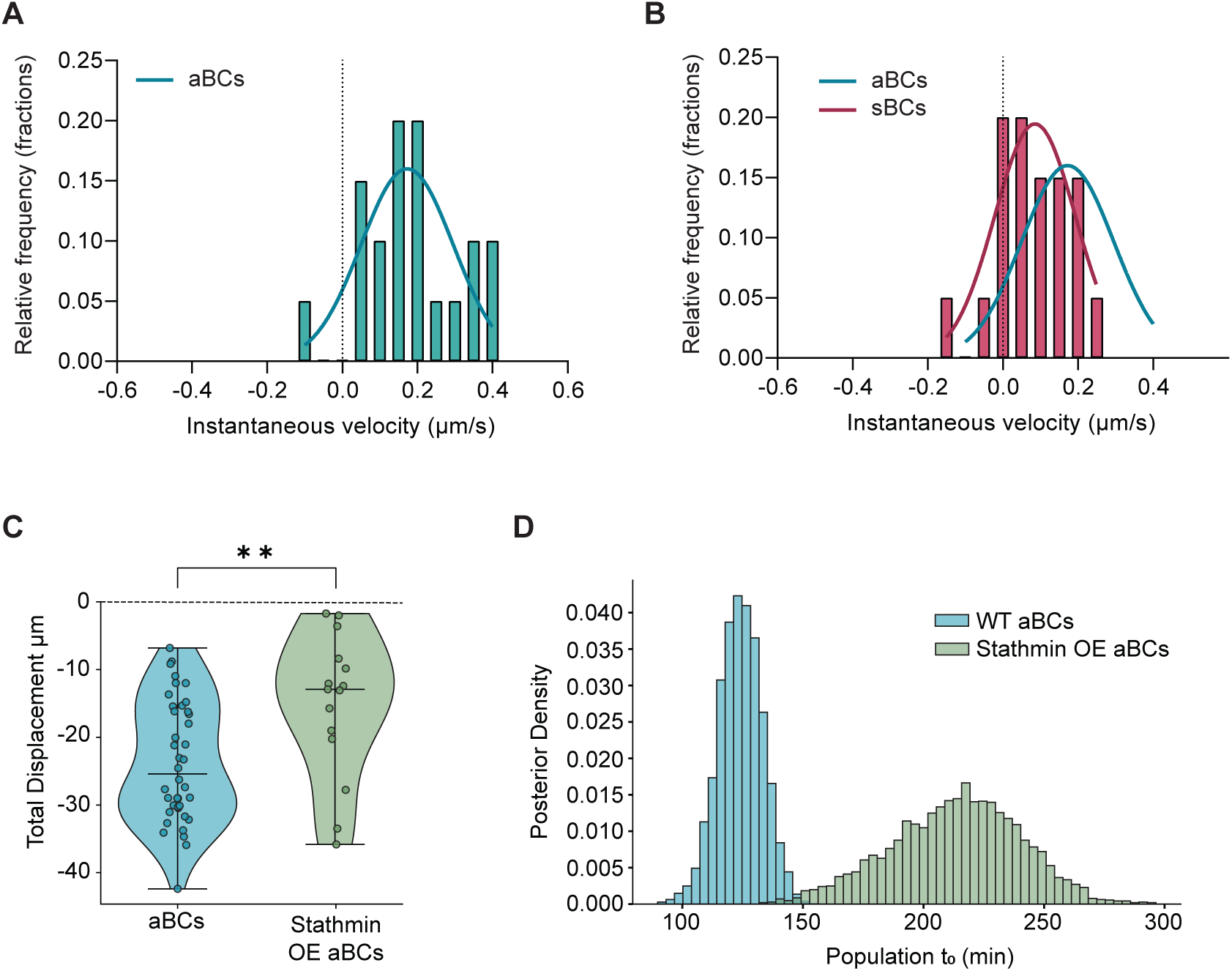
Apically born Bipolar Cells undergo MT-dependent somal translocation. **(A)** Instantaneous velocity distribution of movement apically born Bipolar Cells (aBCs) during first 100 minutes of translocation. Curve shows nonparametric density estimates. **(B)** Instantaneous velocity distribution of movement subapically born BCs (sBCS) during first 100 minutes of translocation. Curves show nonparametric density estimates. **(C)** Violin plots of the distribution of the total displacement of control aBCs and Stathmin overexpressing aBCs over the first 100 minutes of translocation. Each dot represents one cell. The horizontal bar is the median. P value (Mann-Whitney) = 0.010. **(D)** Distribution of population level t_0_ in control aBCs and Stathmin overexpressing aBCs.

**Figure S2.**
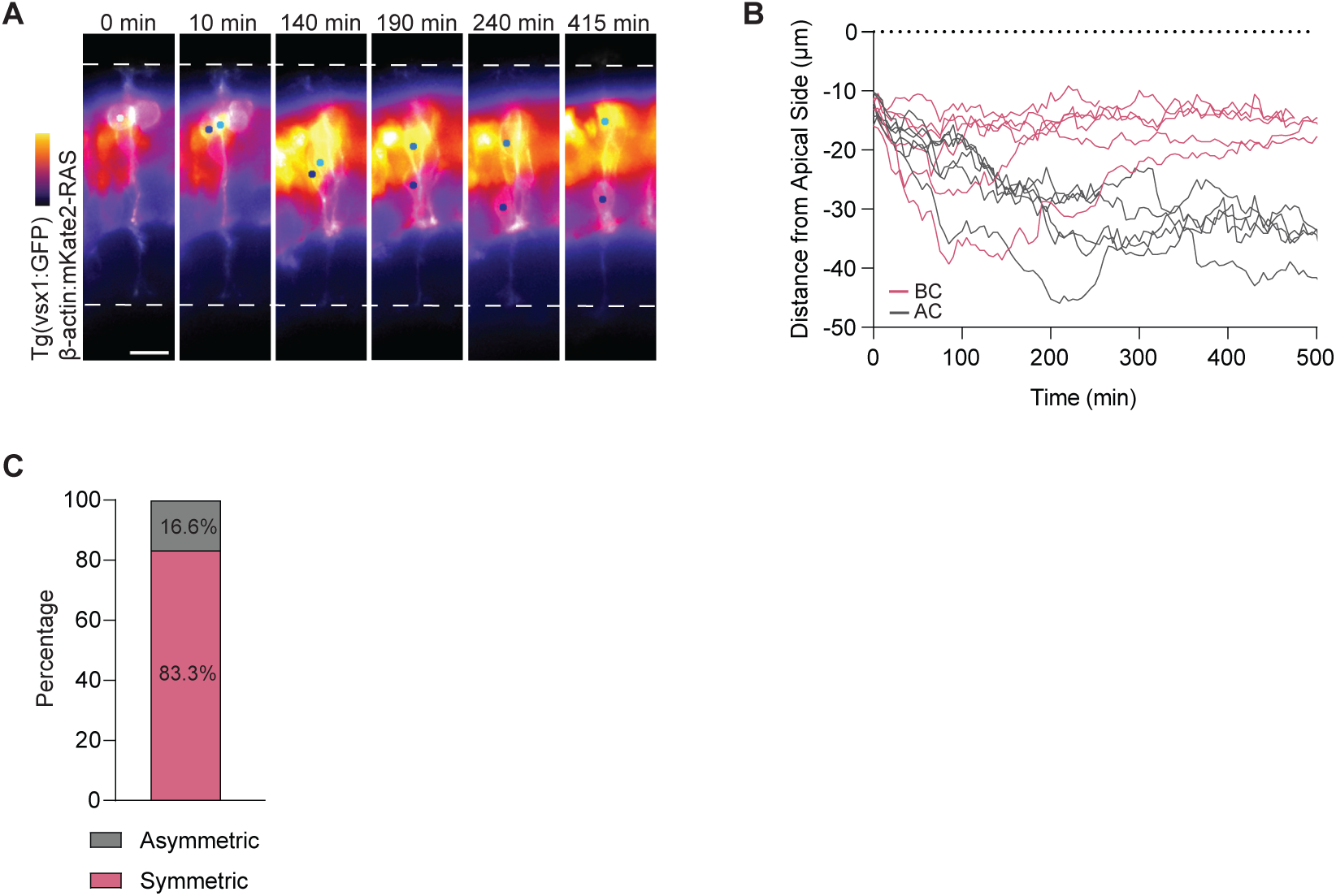
Subapically born Bipolar Cells undergo MT-independent translocation. **(A)** Live imaging of asymmetric BC division and sister cell translocation upon subapical division. Time is shown in minutes. Light blue dot shows BC sister cell and dark blue dot shows Amacrine sister cell followed. Dashed lines represent the apical surface and basal lamina. Scale bar, 10 μm. **(B)** Kinetics of BC-AC sister cell translocation. Time 0 corresponds to the time of division. 10 single trajectories are shown. Dotted line marks the apical surface. (n=10 cells, N= 5 embryos). **(C)** Distribution of asymmetric vs symmetric divisions observed in live imaging experiments.

**Figure S3.**
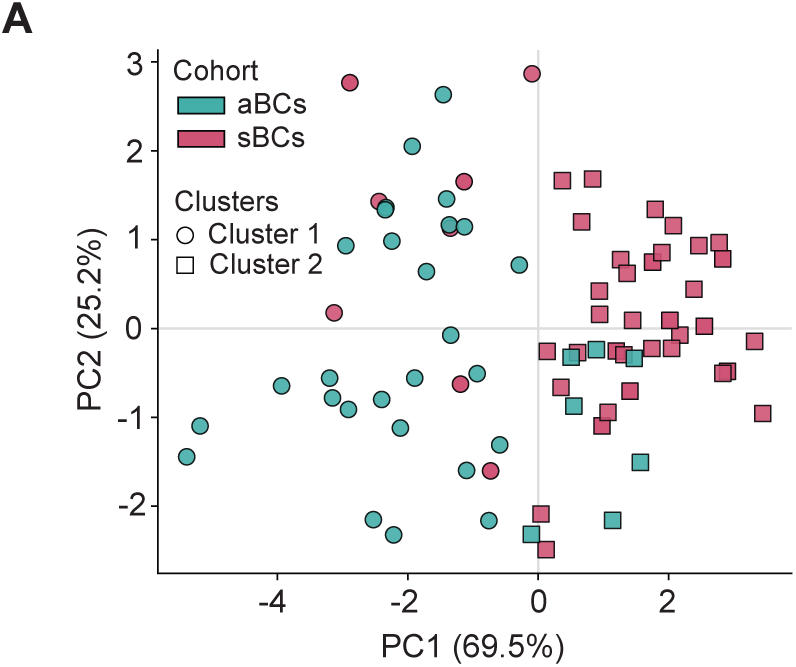
Apically born BCs and subapically born BCs employ different translocation modes. **(A)** Principal Component Analysis (PCA) of kinetic features from apically (cyan) and subapically born BCs (magenta). Symbol shape indicates unsupervised cluster assignment (k=2). Colour indicates cohort. PC1 and PC2 explain 69.5% and 25.2% of total variance, respectively. Cluster separation along PC1 is statistically significant (PERMANOVA p value = 0.0001).

## Video Legends

**Video S1. Apically born Bipolar Cells undergo MT-dependent somal translocation, related to Figure 2**.

**Part A.** Translocation of control apically born Bipolar Cells (aBC). *Tg(vsx1:GFP)* (cyan) labels BCs and *Tg(*β*-actin:mKate2-RAS)* (magenta) labels cell membranes.

**Part B.** Stable microtubule (MT) dynamics is basally translocating aBCs. β-actin:StableMARK-NeonGreen (cyan) labels stable MTs and vsx1:mKate2-RAS (magenta) labels aBCs.

**Part C.** Basal translocation of Stathmin overexpressing aBCs. vsx1:GFP-CAAX (cyan) labels BCs and hsp70:Stathmin1:mKate2 (magenta) labels Stathmin overexpressing cells.

Dots label tracked cells. Time is shown in hours:minutes. Scale bar is 10 μm.

**Video S2. Subapically born Bipolar Cells undergo MT-independent somal translocation, related to Figure 3**.

**Part A.** Translocation of control subapically born Bipolar Cells (sBC). *Tg(vsx1:GFP)* (cyan) labels BCs and *Tg(*β*-actin:mKate2-RAS*) (magenta) labels cell membranes. White arrow indicates presence of apical process. Yellow arrow indicates retraction of apical process.

**Part B.** Stable microtubule (MT) dynamics is basally translocating sBCs. β-actin:StableMARK-NeonGreen (cyan) labels stable MTs and vsx1:mKate2-RAS (magenta) labels sBCs. White arrow indicates presence of apical process. Yellow arrow indicates retraction of apical process. **Part C.** Basal translocation of Stathmin overexpressing sBCs. vsx1:GFP-CAAX (cyan) labels BCs and hsp70:Stathmin1:mKate2 (magenta) labels Stathmin overexpressing cells.

Dots label tracked cells. Time is shown in hours:minutes. Scale bar is 10 μm.

**Video S3. Basal translocation of subapically born Bipolar Cells is driven by collective tissue dynamics, related to Figure 4**.

**Part A.** Translocation of control apically born Bipolar Cell (aBC) in Stathmin overexpressing retina. *Tg(vsx1:GFP)* (cyan) labels BCs and *Tg(hsp70:Stathmin1:mKate2)* (magenta) labels Stathmin overexpressing cells.

**Part B.** Translocation of control subapically born Bipolar Cell (sBC) in Stathmin overexpressing retina. *Tg(vsx1:GFP)* (cyan) labels BCs and *Tg(hsp70:Stathmin1:mKate2)* (magenta) labels Stathmin overexpressing cells.

**Video S4. Disruption of Photoreceptor layer integrity alters Bipolar Cell translocation behaviour at late developmental stages, related to Figure 5**.

Translocation of late-born Bipolar Cell in photoreceptor-depleted are of *prdm1* KO retina. vsx1:GFP-CAAX (cyan) labels BCs and *Tg(ath5:gap-RFP)* (magenta) labels remaining retinal neurons including photoreceptors.

